# Reply to Li et al. “Is *Oculudentavis* a bird or even archosaur?”

**DOI:** 10.1101/2020.06.12.147041

**Authors:** Jingmai O’Connor, Lida Xing, Luis Chiappe, Lars Schmitz, Gang Li, Qiru Yi

## Abstract

We welcome any new interpretation or alternative hypothesis regarding the taxonomic affinity of the enigmatic *Oculudentavis khaungraae*. However, here we demonstrate that Li et al. have failed to provide conclusive evidence for the reidentification of HPG-15-3 as a squamate. We analyse this specimen in a matrix that includes a broad sample of diapsid reptiles and resolve support for this identification only when no avian taxa are included. Regardless of whether this peculiar skull belongs to a tiny bird or to a bizarre new group of lizards, the holotype of *Oculudentavis khaungraae* is a very interesting and unusual specimen, the discovery of which represents an important contribution to palaeontology. Its discovery documents a potential new case of convergent evolution in reptiles, while highlighting the importance of amber deposits for documenting taxa not recorded in sedimentary deposits.

## Introduction

We welcome any new interpretation or alternative hypothesis regarding the taxonomic affinity of the enigmatic *Oculudentavis khaungraae*. Several of the squamate morphologies described by Li et al. were noted by ourselves in the original manuscript (e.g., pleurodont dentition, morphology of the eye)^1^. However, we will argue that other features which Li et al. describe as unusual for archosaurs are not incompatible with our original interpretation.

## Results and Discussion

The antorbital fenestra has been lost multiple times independently for different biomechanical reasons in the Archosauria (Aves, Hadrosaurinae and crocodilians)^2^. For example, the absence of an antorbital fenestra has evolved multiple times within Neornithes (modern birds; Figure 1a), at least once in enantiornithines^2-4^, and a configuration similar to *Oculudentavis* has evolved in some Mesozoic marine crocodyliforms^5^.

**Figure 1.**
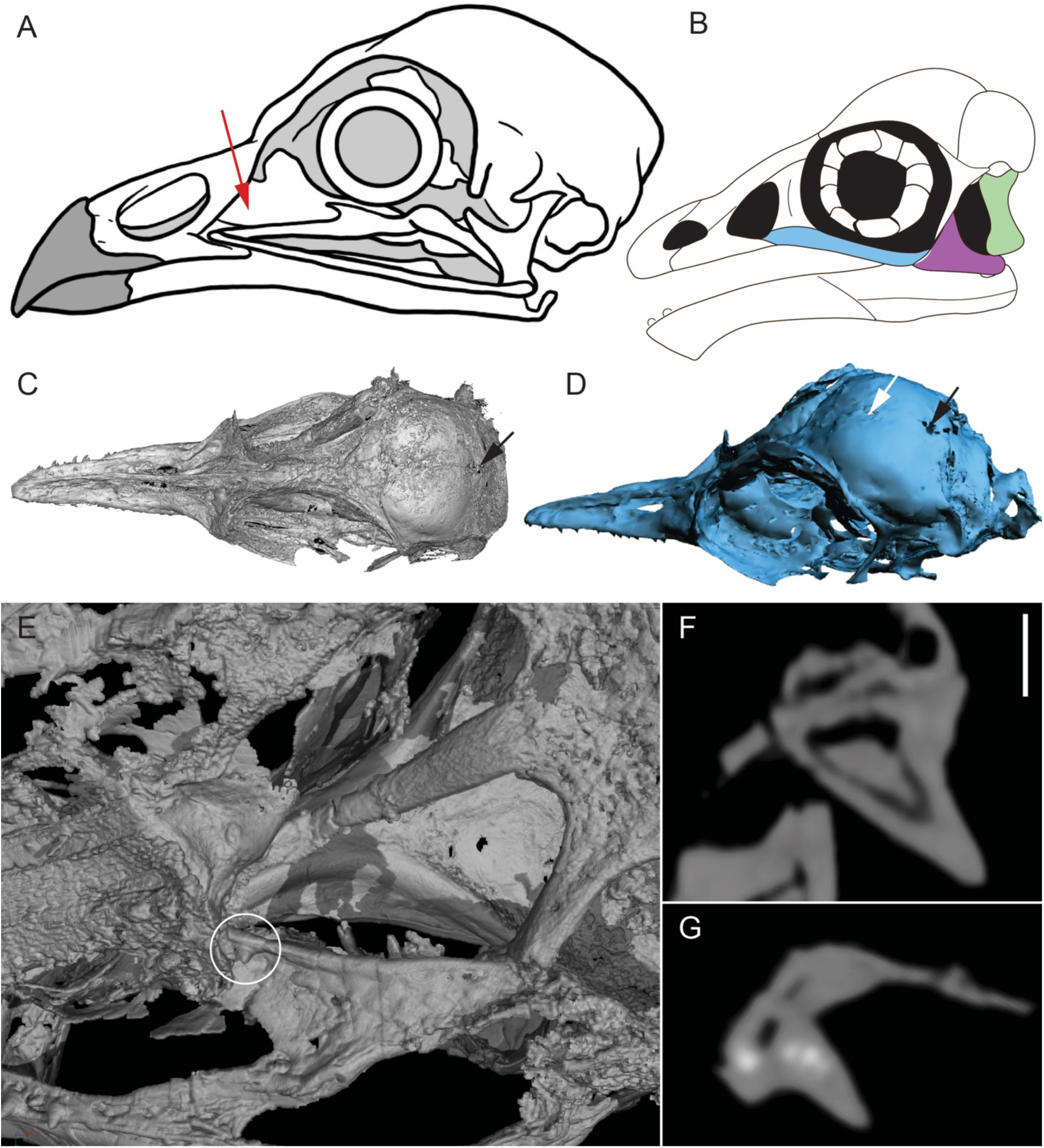
Additional information to clarify arguments related to the phylogenetic affinity of *Oculudentavis* HPG-15-3. (A) skull drawing of a Crested Partridge (*Rollulus rouloul*) – note that the antorbital fenestra (red arrow) and orbit are confluent, a feature that has arisen multiple times in Aves and the Archosauria; (B) reconstruction of *Jeholornis* (originally from^17^) revised to show the contact between the premaxilla and maxilla (colour scheme follows Li et al. – the jugal is blue, the quadratojugal is purple, and the quadrate is green); (C, D) skull of HPG-15-3 indicating position of purported pineal foramen – the white arrow indicates the slight opening Li et al. refer to this feature but we regard as too small to represent the pineal foramen and the black arrows indicate a larger opening that is more consistent with the morphology of this feature excepting the fact it is located too far caudal and is interpreted as poor preservation; (E) view of the palate in HPG-15-3 showing the presence of a tooth-like projection on the right pterygoid (white circle) but not the left; (F) raw CT slice showing the cross section of a maxillary tooth; (G) raw CT slice showing the cross-section of the purported pterygoid tooth – note the white spots indicative of high density that are absent in the maxillary tooth and strongly suggest that this projection on the pterygoid is not a skeletal morphology but an impurity. Scale bar in F also applies to G and equals 20 millimetres.

Furthermore, the jugal in *Oculudentavis* does in fact form the ventral margin of the orbit, but the maxilla extends below it in a long overlapping articulation as far as the tooth row, as in *Ichthyornis*^6^, and therefore this morphology is not without precedent in Aves. Li et al. misinterpret the *Jeholornis* reconstruction in which the premaxilla-maxilla contact is not illustrated – as depicted, the jugal forms the ventral margin of the orbit and the element interpreted by Li et al. as the jugal is in fact the quadratojugal (Figure 1b). The skull of *Jeholornis* has not been described in great detail^7,8^ and the reconstructions in Figure 4^1^, which vary in accuracy based on available information, were meant to be used to compare size and not for detailed anatomical comparisons.

The absence of an infratemporal fenestra cannot be conclusively decided based on the holotype specimen. In the original paper we do not describe a quadratojugal^1^, and as Li et al. themselves point out, the skull is crushed in this region. The quadratojugal is typically small in Cretaceous birds^9^ and rarely preserves, even among the largely complete skeletons collected in the Jehol Biota. Thus, our interpretation that a small quadratojugal was present, enclosing the infratemporal fenestra, and either not visible or not preserved^1^ is reasonable.

A strict dichotomy between pleurodonty in squamates and thecodonty in archosaurs may be an oversimplification^10,11^, despite its irrefutable phylogenetic importance. We noted the pleurodont implantation of the teeth of *Oculudentavis* and, at the time, hypothesized that this unusual feature was related to the interpretation that *Oculudentavis* represented a miniaturized predator as suggested by numerous features^1^. Although there is no precedent for this hypothesis, there are also no miniaturized toothed archosaurs with which to derive analogues. However, the geometry of tooth implantation varies among squamates as well as archosaurs^11^. Within squamates, mosasaurids have thecodont dentitions, while some archosaurs – including the fossil birds *Hesperornis* and juveniles of *Ichthyornis*, and juveniles of the crocodilian *Caiman crocodilus* – have aulacodont dentitions (teeth set in a longitudinal groove without alveoli)^11^. Although rare, deviations from the general pattern within clades are therefore not unheard of, and new discoveries in palaeontology often reveal new morphological combinations not previously seen in the biological record.

Although features like a pineal foramen or pterygoid teeth would certainly and without a doubt remove any reason to hypothesize *Oculudentavis* is avian, we disagree with Li et al. in the presence of these traits. The pineal foramen is a distinct, circular opening on the top of the skull present in many squamates. However, the slight gap indicated by Li et al. is more likely due to disarticulation between the unfused parietals. A more distinct opening is present between the parietals near their caudal margin but this opening is too far caudal to represent the pineal foramen^12^ supporting our original interpretation that this gap is due to poor preservation^1^ (Figure 1c,d). A single tooth-like projection is visible on the right pterygoid, but not the other, questioning whether this is also a true feature. The raw data reveals white spots in the purported pterygoid tooth indicating high density (absent in the premaxillary, maxillary and dentary teeth) consistent with interpretation of this projection as an impurity and not a skeletal feature (Figure 1e-g). This disagreement also highlights the nature of CT data in that the results are sometimes subject to interpretation.

Li et al. criticize our phylogenetic analysis yet provide none themselves. However, this does highlight a weakness of a majority of phylogenetic analyses utilized to describe new taxa. If a new specimen is identified as a bird it is analysed in a matrix targeted at birds; if the specimen is identified as a lizard, it is analysed in a matrix targeted at lizards. Descriptions of new taxa rarely include phylogenetic datasets targeted at higher level relationships such as all of Reptilia or Amniota that would be capable of testing alternative placements. We added *Oculudentavis, Archaeopteryx, Sapeornis, Pterygornis, Rapaxavis*, an unnamed enantiornithine^4^, *Ichthyornis, Parahesperornis*, and *Llallawavis* into the recent Pritchard and Sues^13^ matrix which broadly samples Diapsida and previously included no avians. Characters related to uncertain morphologies such as the presence of a quadratojugal and postfrontal were scored with “?”.

Our parsimony-based phylogenetic analysis run using TNT^14^ placed *Oculudentavis* in Aves (Fig. 2). Forcing a relationship with squamates required 10 additional steps. Even when *Archaeopteryx* was the only included avian *Oculudentavis* was resolved as a bird and forcing it to fall among squamates required three additional steps and produced deep polytomies. However, removal of all avian taxa results in *Oculudentavis* being resolved among squamates.

**Figure 2.**
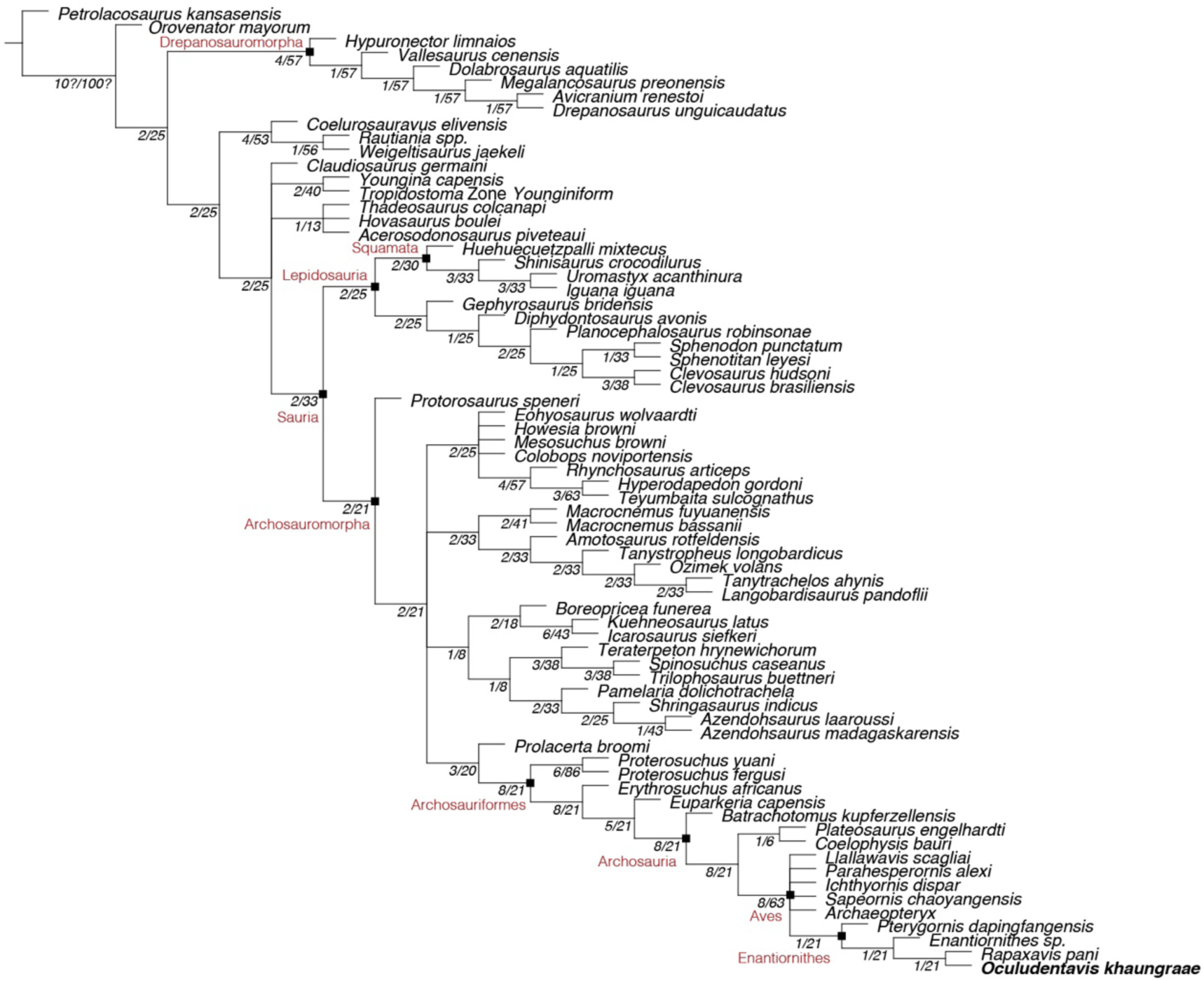
Phylogenetic position of *Oculudentavis* using the Pritchard & Sues (2019) matrix. In the strict consensus tree of 29 retained trees, *Oculudentavis* is resolved nested among enantiornithines (length = 1381; consistency index = 0.265; retention index = 0.632). Numbers indicate absolute and relative Bremer supports. Constraining *Oculudentavis* to Squamata requires ten additional steps. Even if only *Archaeopteryx* is included, *Oculudentavis* is resolved as an avian although removal of all birds results in the placement of *Oculudentavis* among Squamata.

As clearly stated in the etymology^1^, the name *Oculudentavis* was not derived to mean a bird with bird-like tooth and eye morphology as described by Li et al., but rather a bird with a toothed eye, referring to the dentition below the eye that also occurs, although less extensively, in *Ichthyornis*^6^. Although in the future new information may prove we are incorrect in our original interpretation and *Oculudentavis* may be added to the list of taxa whose names have become misnomers (e.g., *Oviraptor, Piscivorenantiornis*)^15,16^, this is in no way due to gross negligence, but rather due to the fact that *Oculudentavis* may represent an outstanding case of convergent evolution between squamates and birds, the likes of which biologists have rarely seen before. Li et al. do not provide conclusive support for their alternative interpretation of *Oculudenatvis*, and their hypothesis is only supported in the absence of avians in the phylogenetically broader cladistic analysis presented here (Fig. 2). However, regardless of whether this peculiar skull belongs to a tiny bird or to a bizarre new group of lizards, the holotype of *Oculudentavis khaungraae* is a very interesting and unusual specimen, the discovery of which represents an important contribution to palaeontology^1^. Its discovery documents a potential new case of convergent evolution in reptiles, while highlighting the importance of amber deposits for documenting taxa not recorded in sedimentary deposits.

## Supporting information

nexus file for phylogenetic analysis

## Author contributions

J.O’C., L.X., L.C., and L.S. designed the response, J.O’C., L.C., and L.S. wrote the response, J.O’C. scored the birds into the matrix, L.S. analysed the matrix, and J.O’C., G.L., L.X., and Q.Y. re-examined the original data.

## Competing Interests statement

The authors declare no competing interests.

## Notes

https://ea-boyerlab-morphosource-01.oit.duke.edu/Detail/SpecimenDetail/Show/specimen_id/30882

https://www.biorxiv.org/content/biorxiv/early/2020/03/18/2020.03.16.993949.full.pdf

### Competing Interest Statement

The authors have declared no competing interest.

